# WGT: Tools and algorithms for recognizing, visualizing and generating Wheeler graphs

**DOI:** 10.1101/2022.10.15.512390

**Authors:** Kuan-Hao Chao, Pei-Wei Chen, Sanjit A. Seshia, Ben Langmead

**Affiliations:** Department of Computer Science, Johns Hopkins University; Department of Electrical Engineering and Computer Sciences, University of California, Berkeley

**Author notes:** These authors contributed equally to this work (Kuan-Hao Chao, Pei-Wei Chen).

## Abstract

**Summary:** A Wheeler graph represents a collection of strings in a way that is particularly easy to index and query. Such a graph is a practical choice for representing a graph-shaped pangenome, and it is the foundation for current graph-based pangenome indexes. However, there are no practical tools to visualize or to check graphs that may have the Wheeler properties. Here we present `Wheelie`, an algorithm that combines a *renaming heuristic* with a permutation solver (`Wheelie-PR`) or a Satisfiability Modulo Theory (SMT) solver (`Wheelie-SMT`) to check whether a given graph has the Wheeler properties, a problem that is NP complete in general. `Wheelie` can check a variety of random and real-world graphs in far less time than any algorithm proposed to date. It can check a graph with 1,000s of nodes in seconds. We implement these algorithms together with complementary visualization tools in the WGT toolkit, available as open source software at https://github.com/Kuanhao-Chao/Wheeler_Graph_Toolkit.

## 1 Introduction

A Wheeler graph is a class of directed, edge-labeled graph that is particularly easy to index and query. It is a generalization of the Burrows-Wheeler-Transform-based FM Index [1], and partly forms the basis for existing pangenome alignment tools such as vg [2, 3].

A graph is a Wheeler Graph when its nodes can be totally ordered according to the co-lexicographical order of the sets of strings spelled out on all paths leading into the nodes. Formally: an edge-labeled, directed graph is a Wheeler graph if and only if there exists a total ordering over its nodes such that 0-indegree nodes come before all other nodes in the ordering, and for all pairs of edges (*u, v*) and (*u′, v′*) labeled *a* and *a′* respectively: (i) *a ≺ a′ → v < v′*, and (ii) *a* = *a*′ ∧ *u* < *u*′ → *v* ≤ *v*′.

Wheeler graphs generalize other graph- and tree-shaped structures important in genomics, including tries, De Bruijn graphs, and the reverse deterministic graphs that can be constructed from multiple sequence alignments [4]. The reverse deterministic graph in this paper refers to the graph defined in Section 6.1 in the 2014 paper proposed by Siren et al [5]. These special cases of Wheeler graphs have been applied in the context of pangenome alignment tools like GCSA [5], HISAT2 [6], and VARI [7].

Despite their popularity, there are no libraries that make it easy to use Wheeler graphs or to check if a particular graph has the requisite properties. This problem is NP-complete in general and hard to approximate [8]. An exponential-time algorithm was proposed in [8], but no implementation is available.

We present WGT, an open source suite for generating, recognizing, and visualizing Wheeler graphs. WGT includes functionality for generating graphs that do or do not have the Wheeler properties. Two generators produce De Bruijn graphs and tries derived from one or more input sequences provided as FASTA. Another generator produces reverse deterministic graphs [5] from multiple sequence alignments. A fourth generator produces random graphs parameterized by the desired number of nodes, edges, distinct edge labels (i.e. alphabet size), and the most number of outgoing same-label edges.

Central to WGT is the fast `Wheelie` algorithm for Wheeler graph recognition. The algorithm combines a *renaming heuristic* with two alternate solvers, both capable of reaching exact solutions to the recognition problem. One solver uses an exhaustive search over possible node permutations, and the other uses a Satisfiability Modulo Theory (SMT) solver [9]. We call the overall algorithm “`Wheelie`”, while we use the names “`Wheelie-Pr`” and “`Wheelie-SMT`” for the versions that use the permutation and SMT solvers respectively. When run on a Wheeler graph, `Wheelie` also reports a node ordering for which the properties are satisfied and indexes the graph into *O, I*, and *L* three bitarrays [4], which are useful inputs to a downstream tool for pattern matching.

Here we benchmark `Wheelie`’s solvers in comparison to each other and to the algorithm proposed by Gibney and Thankachan [8]. We benchmark with a variety of input graphs, including graphs derived from real multiple alignments of DNA and protein sequences. We also use randomly-generated graphs with various configurable characteristics. Finally, we implement and demonstrate a visualizer that allows the user to picture the graph in light of the Wheeler properties.

In the following, *G* denotes a directed graph, *N* its set of nodes, and *E* its set of edges, with *n* = |*N* | and *e* = |*E*|. Σ denotes the set of edge labels appearing on at least one edge, with *σ* = |Σ|.

## 2 Results

Graphs used for evaluation were generated using WGT’s generator algorithms, which can produce (a) De Bruijn graphs, (b) tries, (c) a reverse deterministic graphs derived from a multiple alignments, (d) *complete* random Wheeler graphs, and (e) a *d-NFA* random Wheeler graphs. All are discussed further in Methods 3.3 and Appendix 2. For graphs that start from biological sequences, we selected 25 genes and downloaded their DNA and protein ortholog alignments in FASTA format from the Esnbmbl Comparative Genomics page [10]. All the experiments are conducted on a 24-core, 48-thread Intel(R) Xeon(R) Gold 6248R Linux computer with 1024 GB memory, using a single thread of execution.

### 2.1 Comparing Wheelie with Gibney & Thanckachan

Gibney and Thankachan’s recognition algorithm [8] (henceforth “G & T”) works by enumerating all possible values for the *O, I* and *L* arrays making up the Wheeler Graph structure as described by Gagie et al [4]. The *O* bitarray is a concatenation of unary codes describing the outdegrees of each node. *I* is a similar bitarray that does the same for indegrees. *L* is a sequence of characters labeling the edges in the order they appear in the *O* array. Further, the inner loop of the algorithm must check if a given assignment for *I, O*, and *L* is isomorphic to the input graph provided.

While the G & T algorithm explores an exponential-sized space, `Wheelie` explores the factorial-sized space of node permutations. While this could represent a disadvantage for `Wheelie`, we hypothesized that `Wheelie` could be made faster with the help of strategies for pruning the search space. `Wheelie` prunes its search by assigning labels to nodes according to their *rough* positions in the order, a strategy we call the “*renaming heuristic*”. This allows `Wheelie` to arrive rapidly at a rough ordering that either (a) reveals a conflict that prevents the graph from having the Wheeler properties, or (b) reduces the problem size for the downstream solving algorithm. The full algorithm is described in Methods 3.1, 3.2. Here we use a version of the algorithm called `Wheelie-Pr`, which begins with the *renaming heuristic* then resolves remaining ambiguities by exhaustively searching over the remaining node permutations. Unlike G & T, the `Wheelie-Pr` algorithm does not need to compute graph isomorphsims.

We conducted a 30-second timeout test on both algorithms using graphs generated from four generators (3.3) including both Wheeler and non-Wheeler graphs. Rather than implement G & T’s entire algorithm, we implemented the enumeration of the *I, O* and *L* arrays but omitted the graph isomorphism check in the inner loop. We reasoned that if `Wheelie-Pr` was faster than the G & T algorithm without the (rather complex) isomorphism check, it would also be faster than the full G & T algorithm. To compare the algorithms, we configured both to perform an exhaustive search, without the possibility of early stopping if a solution is found. This differs from `Wheelie-Pr`’s default behavior, which allows it to stop upon finding a node ordering for which the Wheeler properties are satisfied. Early stopping is still possible for `Wheelie-Pr` in these experiments, since it might identify a conflict that proves the graph is non-Wheeler. We took 25 multiple orthologue alignments, both their DNA and amino acid (AA) sequences, and extracted the first 4 rows of each. To reduce graph size, we truncated the graphs with respect to the multiple-alignment columns. We tested on three types of graphs, De Bruijn graphs, tries, and random Wheeler graphs, that are known Wheeler graphs, and two types of graphs, pseudo-De Bruijn graphs and reverse deterministic graphs, that are not guaranteed Wheeler graphs. Pseudo-De Bruijn graphs are graphs where the nodes correspond to k-1-mers in the multiple alignment, but where we do not collapse identical k-1-mers into single nodes.

For De Bruijn graphs and pseudo-De Bruijn graphs, we took columns 1 to 200 and set *k* to 3 to 9; for tries, we took columns 1 to 200; for reverse deterministic graphs, we took columns 2 to 41. We also benchmarked with a series of randomly generated graphs with *n* set to 3 to 33, *e* from 3 to *n* and *σ* from 1 to 21. The number of each type of graphs and their node and edge numbers are shown in Figure 1B, and the arguments of each generator can be found in the WGT Github repository: https://github.com/Kuanhao-Chao/Wheeler_Graph_Toolkit.

**Figure 1:**
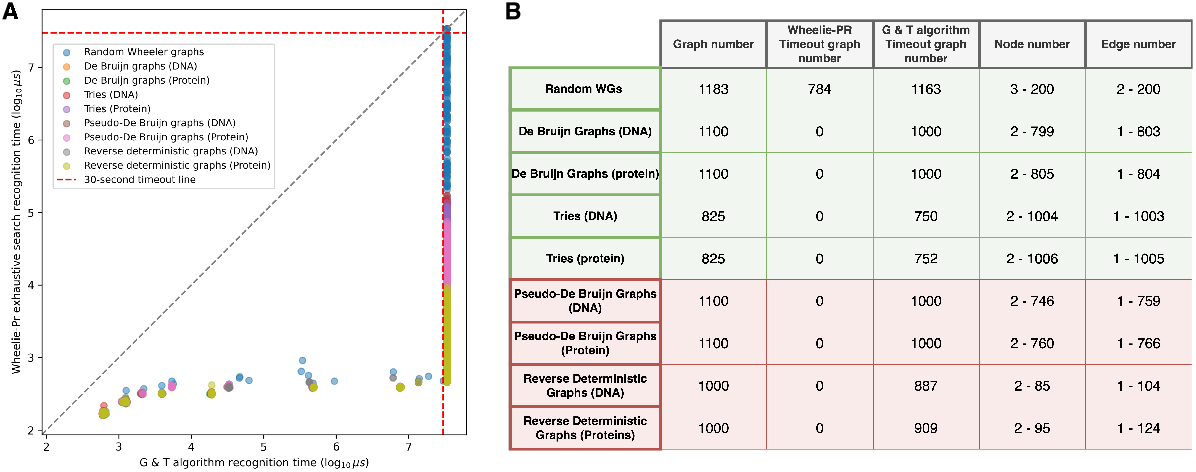
(A) Recognition time comparison between `Wheelie-Pr` exhaustive search and the G & T algorithm using (1) De Bruijn graphs, (2) tries, (3) pseudo-De Bruijn graphs and (4) reverse deterministic graphs generated from DNA and protein alignments, and (5) random graphs generated with given *n, e* and *σ*. `Wheelie-Pr` recognition time is on the y-axis, and G & T recognition time on the x-axis, both on a log_10_ microsecond scale. Each dot represents a graph. Dots beyond the red lines denote inputs for which the tool timed out after 30 seconds. (B) The graph number, timeout graph number of `Wheelie-Pr` and the G & T algorithm, and ranges of node and edge number of each type of graphs. Rows in green describe graphs that are guaranteed Wheeler. Rows in red describe graphs that are not guaranteed to be Wheeler; i.e. some instances are Wheeler and some are not.

Figure 1A shows that `Wheelie-Pr` is significantly faster, allowing it to recognize a range of Wheeler and non-Wheeler graphs. `Wheelie-Pr` runtimes generally range from 100-1,000 microseconds, with 784 random-graph inputs causing `Wheelie-Pr` to time out. In sum, the only type of graph that caused `Wheelie-Pr` to time out is the random graph whereas 8,461 graphs distributed in all types of benchmarked graphs caused G & T to time out.

### 2.2 Visualizing and characterizing challenging graphs

We selected a De Bruijn graph with edges being k-mers and nodes being k-1-mers where *k* = 4 from the Figure1 benchmarks. This graph was derived from the first four rows of the multiple alignment of STAU2 DNA orthologues with sequence length 4. We first visualized it using Graphviz [11] (Figure 2A). We ran `Wheelie-Pr` to find an ordering for which the Wheeler properties hold (Figure 2B). Finally, we visualized the graph using WGT’s Python-based visualizer, which draws the ordered nodes in two replicas, with outgoing edges leaving one replica (Figure 3, top rows) and entering the other (bottom rows). For a valid Wheeler ordering, nodes with no incoming edges will appear leftmost, nodes with incoming edges of the smallest character will come next, nodes with incoming edges of the next-smallest character next, etc. Further, no two same-color edges will cross each other. In this way, the diagram, first described by Boucher et al [12] – makes it visually obvious when an ordering has yielded the Wheeler properties.

**Figure 2:**
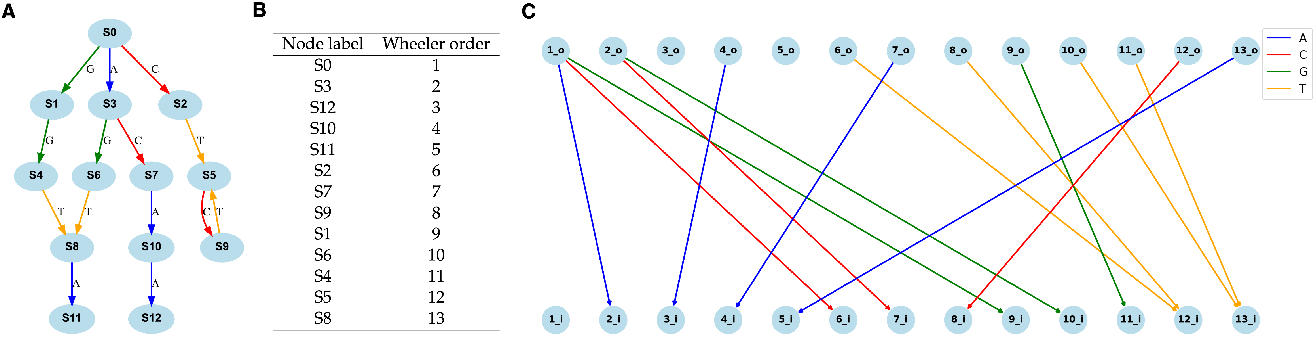
(A) A *k* = 4 De Bruijn graph outputted from WGT’s De Bruijn graph generator. It is the visualization from Graphviz online visualizer. (B) The recognition result showing the Wheeler ordering outputted from `Wheelie-Pr`. (C) The output from WGT’s visualizer. Nodes are duplicated into two rows ordered in Wheeler ordering.

**Figure 3:**
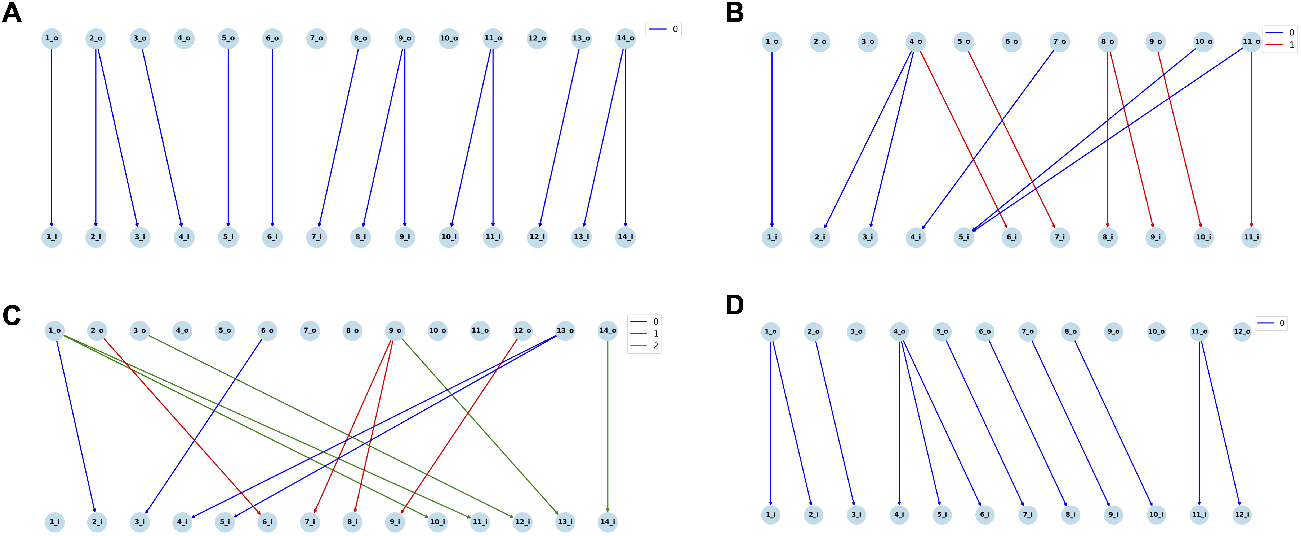
Examples of (A) 2-NFA with 1 label (B) 2-NFA with 2 labels (C) 2-NFA with 3 labels (D) 3-NFA with 1 label, from outlier random graphs (blue dots) in Figure1.

We sought to understand which graphs require the most time for recognition. After investigated the “outlier” graphs where Wheelie-PR timed out with these tools, we found that the graphs requiring the most recognition time tended to have nodes with many outgoing same-label edges. Following Alanko et al [13], we use the term ≤ *d*-NFA to describe a Wheeler Graph where all nodes have ≤ *d* outgoing same-label edges, and at least one node has exactly *d* outgoing same-label edges. The De Bruijn graph shown in Figure 2 is a 1-NFA. Figure 3 A-C are 2-NFAs with *σ* equal to 1, 2 and 3 respectively. Figure 3 D is a 3-NFA with *σ* = 1. Note that the *d ≥* 5 case is the one proven to be NP-complete [8]. De Bruijn graphs and tries are 1-NFAs.

### 2.3 Recognizing challenging graphs with Wheelie-SMT

Motivated by previous work that showed how boolean satisfiability formulations can solve special cases of the recognition problem [13], we hypothesized that Satisfiability Modulo Theories (SMT) solvers [9] could solve all or part of the Wheeler-graph recognition problem. SMT has found many uses in artificial intelligence and formal methods for hardware and software development. As a generalization of the Boolean Satisfiability (SAT) [14], SMT allows us to encode the Wheeler graph properties in a fairly straightforward way, building from the propositional logic formulas in the definition.

We conducted two series of 1,000-second timeout tests using graphs generated from the random generator comparing (1) `Wheelie-Pr` (*renaming heuristic* plus permutation) versus (2) `Wheelie-SMT` (*renaming heuristic* plus SMT) on different types of *d*-NFA (2.3.1) and various sizes of random graphs (2.3.2).

#### 2.3.1 Recognizing *d*-NFAs

We fixed *n* = 1000, *e* = 3000 and *σ* = 4 and randomly generated *d*-NFAs with *d* from 1 to 8 and each group with 20 graphs. Figure4 shows that both solvers can solve graphs swiftly when *d* is 1 and 2; as *d* grows beyond 2, all tools require much more time, demonstrating that *d* impacts the hardness of recognition problem in practice. `Wheelie-SMT` outperforms `Wheelie-Pr` and avoids any timeouts; `Wheelie-Pr` has some timeouts starting at *d* = 3 (4 out of 20 graphs), and consistently times out when *d ≥* 4.

**Figure 4:**
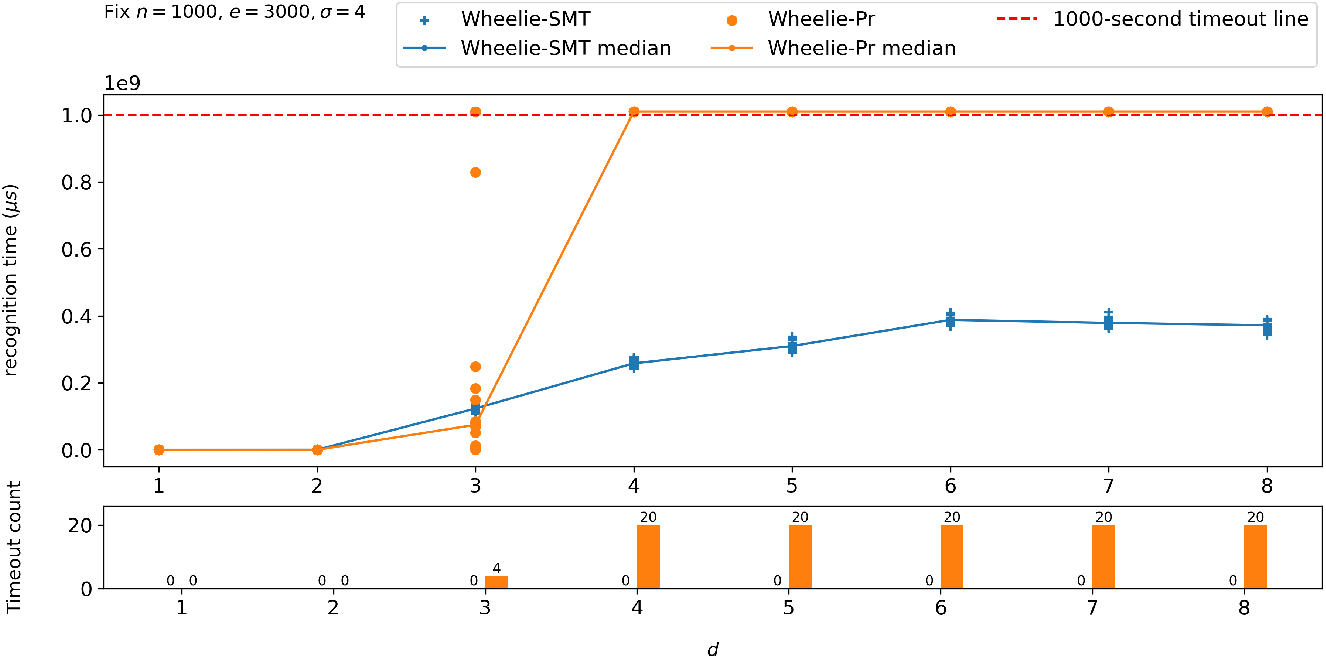
Recognition time for `Wheelie-SMT` and `Wheelie-Pr` as a function of the *d* parameter of the *d*-NFA. Upper panel plots recognition time versus *d* and includes a line connecting the medians. 20 graphs were tested for each *d*. The bottom bar chart shows the number of timeouts.

Further, we observed that when *d ≥* 6, the median curve for `Wheelie-SMT` plateaus. This is because *n* and *e* are too small for the *d*-NFA generator to produce uniformly distributed *d*-NFAs under the given parameters. More precisely speaking, the hardness of the recognition problem is a function of the distribution of nodes having *d −* 1, *d −* 2, …, 1 out-going edges with the same labels. As an example, take a *d*-NFA *G* that has one node with *d* same-label out-going edges, and the rest of the nodes having at most one outgoing same-label edge. Recognizing *G* is not harder than recognizing a uniformly-distributed *d −* 1-NFA. In short, we observed that higher *d*s generally led to a harder recognition problem, but the true level of hardness was also a function of *n, e* and *σ*.

#### 2.3.2 Recognizing random Wheeler graphs

We defined “graph size” as *n* and “label density” as *e/σ*. We then benchmarked various sizes of random graphs while varying these parameters.

We first fixed the number of edges (*e* = 8, 000) and labels (*σ* = 4) while scaling graph size *n* from 2, 000 to 8, 000. Figure5A shows that as *n* grows, `Wheelie-SMT` outperforms `Wheelie-Pr` significantly. `Wheelie-Pr` starts to time out in some cases when *n* = 2, 000, and most cases when *n ≥* 2, 500. In contrast, `Wheelie-SMT` can solve all cases with *n* up to 4, 000, and most cases when *n* = 4, 500.

**Figure 5:**
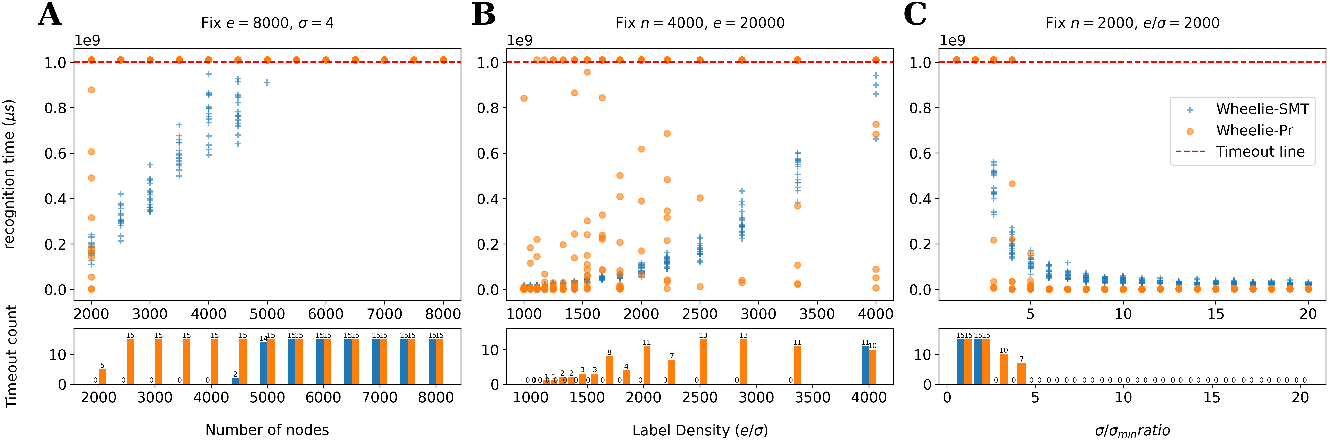
Recognition time comparison for `Wheelie-SMT` and `Wheelie-Pr` for various random Wheeler graphs. Three experiments were conducted. (A) Experiment 1: fixing *e* = 8, 000, *σ* = 4 and scaling up the graph size (*n* from 2, 000 to 8, 000). (B) Experiment 2: fixing *n* = 4, 000, *e* = 20, 000 and scaling up the label density (*e/σ* from 1, 000 to 4, 000). (C) Experiment 3: Fixing both graph size (*n*) and label density (*e/σ*), and scaling up both *e* and *σ* (*σ/σ*_*min*_ from 1 to 20). The upper-panel plots show the recognition time versus the scale up parameter in microsecond scale. Each dot represents a graph. Dots beyond the red dashed line means timeouts. Plots in the lower panel are the timeout count bar charts.

We then fixed the “graph size” (*n* = 4, 000) and number of edges (*e* = 20, 000) and varied the “label density” (*e/σ* from 1, 000 to 4, 000). Figure 5B shows that as the label density increases, the graphs take more time to solve. Comparing `Wheelie-Pr` and `Wheelie-SMT`, we can see that there are more timeout cases in `Wheelie-Pr` from 1, 200 to 4, 000 (most are timeouts when *e/σ ≥* 2, 500) whereas the timeout cases only occur in `Wheelie-SMT` when label density is 4, 000.

In a third experiment, we fixed the graph size (*n* = 2, 000) and varied the number of edges (*e*) and labels (*σ*) while fixing the label density ratio (*e/σ* = 2, 000). Figure5C shows that as more edges and labels are added, the recognition problem becomes easier. In short, this is because adding more constraints to *G* breaks more of the ties that would otherwise obstruct `Wheelie`’s *renaming heuristic*. Comparing `Wheelie-Pr` to `Wheelie-SMT`, Figure5C shows that the solvers perform similarly, with `Wheelie-Pr` performing slightly better when *σ/σ*_*min*_ ratio gets larger (≥ 5). These are likely cases where the graph is sufficiently easy to recognize that the overhead of setting up the SMT setup problem becomes harmful. When *σ/σ*_*min*_ gets smaller (3 and 4), `Wheelie-SMT` is able to solve all 15 cases, whereas `Wheelie-Pr`’s times out for about half the cases.

### 2.4 Benchmarking Wheelie-SMT alone

To isolate the effect of the `Wheelie` *renaming heuristic*, we conducted a 30-second timeout test with 60 seconds timeout penalties on (1) `Wheelie-SMT` (*renaming heuristic* plus SMT) and (2) a pure SMT solver starting from scratch, without the constraints it would otherwise receive from the *renaming heuristic*. We bench-marked these using two generators from DNA alignments: (1) De Bruijn graphs generated with options `-k` from 5 to 8, `-l` from 100 to 2, 000 and `-a` from 6 to 10, 225 graphs in total and (2) reverse deterministic graphs generated with options `-l` from 100 to 500 and `-a` from 4 to 6, in total 225 graphs.

Figure6 shows cactus plots on De Bruijn graphs and reverse deterministic graphs. Cactus plot is an aggregated sorted time plots widely used in solver competitions. It shows how many problems a solver can solve in a limited time period. In Figure6A, `Wheelie-SMT` solved the whole De Bruijn graph set in around 6.5 seconds whereas the pure SMT approach solved it in around 820 seconds. For reverse deterministic graphs (Figure6B), `Wheelie-SMT` solved the whole set in less than 9 seconds whereas the pure SMT approach solved it in around 10,170 seconds.

**Figure 6:**
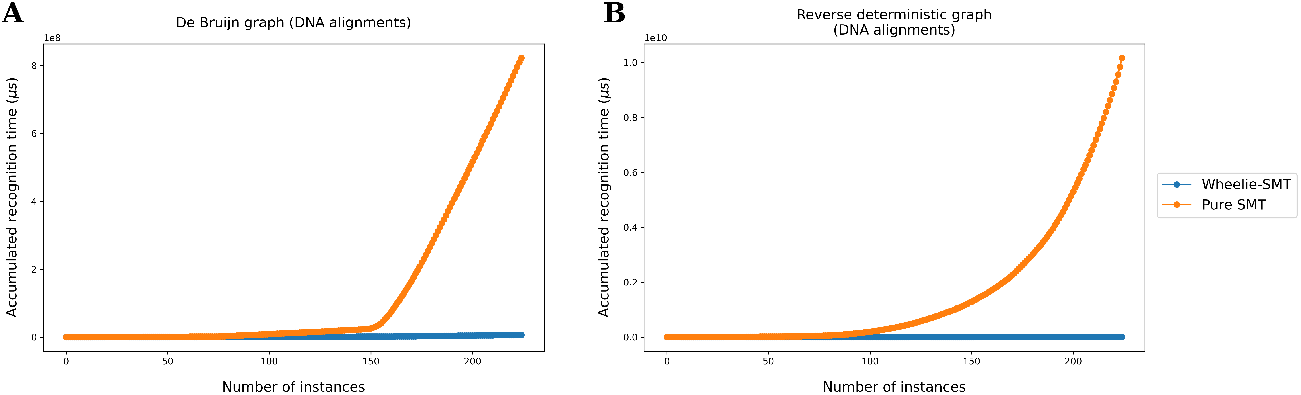
The cactus survival plots of (A) De Bruijn graphs and (B) reverse deterministic graphs generated from DNA alignments using WGT’s generators. They show the aggregated time comparison between `Wheelie-SMT` and pure SMT without the constraints from the *renaming heuristic*..

We concluded that the *renaming heuristic* is a crucial step, since it greatly narrows the space of possible node ordering that must be resolved by the SMT solver. `Wheelie-SMT` can solve graphs several orders of magnitude larger than a pure SMT approach.

## 3 Methods

### 3.1 Wheelie and the *renaming heuristic*

`Wheelie` explores the space of possible node orderings until arriving either at a conflict (e.g. a node with distinctly labeled incoming edges) or an ordering for which the Wheeler properties hold. While this is a large (*n*!-sized) search space, `Wheelie` prunes the space by assigning labels to nodes according to their *rough* position in the overall order. Initially, a rough ordering is determined according to the labels of the immediate incoming edges for each node, following the Wheeler requirement that *a ≺ a′ → v < v*^*′*^ for all edge pairs. This rough ordering is refined over the course of a procedure that iterates either until the rough ordering becomes total ordering, or until the rough ordering stabilizes. In the latter case, the remaining ambiguities are resolved by a non-heuristic solver. This procedure is detailed in Algorithm 1 and illustrated in Figure 7.

**Figure 7:**
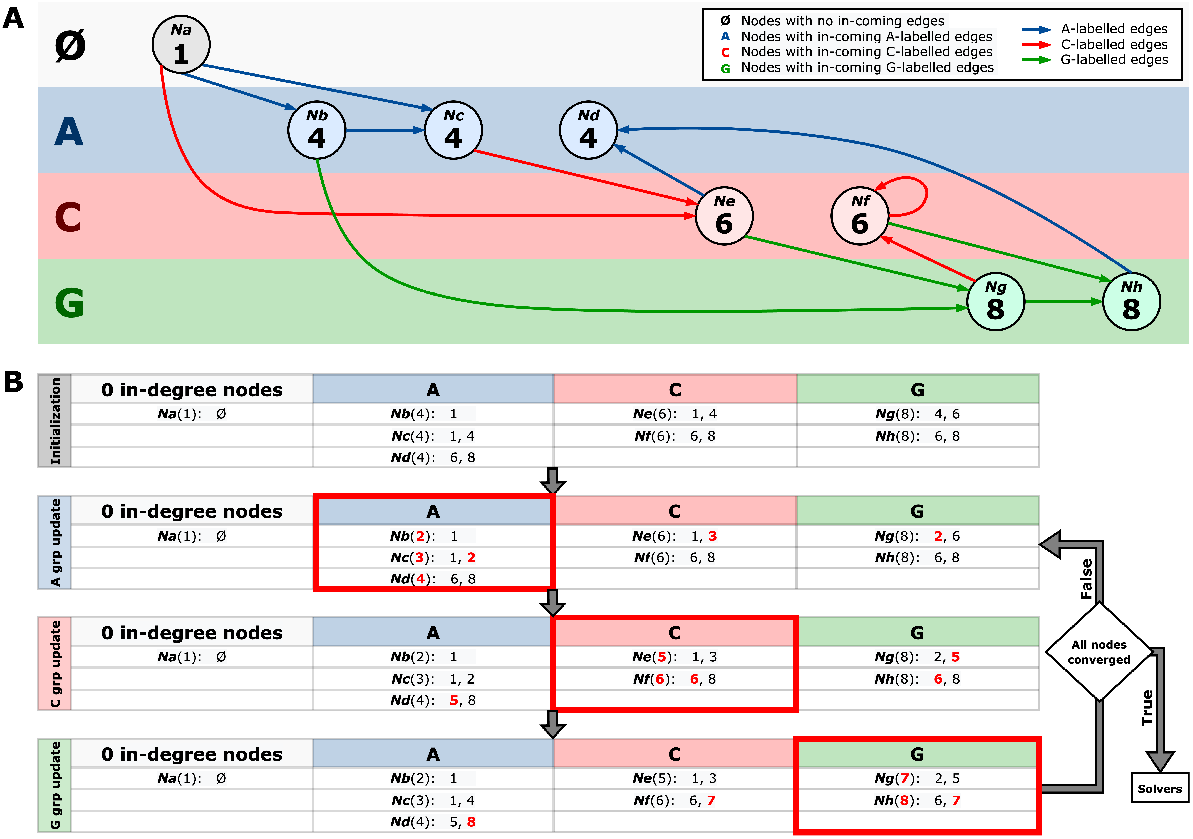
Illustration of the *renaming heuristic*. (A) an 8-node graph with nodes divided into four groups according to in-coming edge label (with Ø representing 0-indegree nodes). (B) presents the workflow of the *renaming heuristic*. The first table in (B) shows the initialized in-node lists for eight nodes. After initialization, the algorithm sorts and relabels nodes in each group until convergence. Then, it passes the range information to either `Wheelie-Pr` or `Wheelie-SMT`.

As the *renaming heuristic* iterates, it repeatedly visits the nodes in groupings according to the label of their incoming edge(s). For each of these groupings, it sorts the edges by sources and destinations in every label group, requiring *O*(II_*g ∈* σ_ *e*(*g*) log_2_ *e*(*g*)) time, where *e*(*g*) is the number of edges labelled as *g*. We observed that many non-Wheeler graphs can be recognized as such directly by the *renaming heuristic*, without requiring a downstream solver.

At each iteration, the algorithm gathers a list of sorted unique temporary orders of nodes that go into it, which we term the “in-node list.” By the Wheeler graph property that requires all edge pairs to satisfy *a* = *a′ ∧ u < u′ → v ≤ v′*, we can find rough orders by sorting the nodes by their in-node lists. Once this has been done for each node group, we reach the end of the current iteration and we check if the rough order changed since the previous iteration. If not, then we say the algorithm has converged and forward any remaining ambiguities to the downstream solver as necessary.

We note that there are similarities between the renaming heuristic and the “forward algorithm” of Alanko et al [15] (Algorithm 2 in that paper). While our renaming heuristic performs an explicit sort within each of its rough grouping, the forward algorithm of Alanko et al uses a pair of nested loops over alphabet characters to visit and partition the nodes in a way that maintains their sorted order implicitly. We discuss this relationship further in Discussion 4 below.

`Wheelie` contains two solvers: `Wheelie-PR` and `Wheelie-SMT`. `Wheelie-PR` takes the output from the renaming heuristic and, for any remaining ties in the ordering, simply tries all possible permutations among the tied nodes. The `Wheelie-SMT` solver is explained in the next subsection 3.2.

### 3.2 Satisfiability Modulo Theories (SMT) solver

Motivated by the use of boolean satisfiability formulations to solve special cases of the recognition problem [13], we hypothesized that Satisfiability Modulo Theory (SMT) solvers [9] could be used to solve all or part of the Wheeler-graph recognition problem. SMT has found many uses in artificial intelligence and formal methods for hardware and software development. As a generalization of the Boolean Satisfiability (SAT) [14], SMT allows us to encode the Wheeler graph properties in a fairly straightforward way, building from the propositional logic formulas in the definition.

An SMT problem decides the satisfiability of a first-order formula with respect to one or more background theories. A *formula* is a set of atoms connected by Boolean connectives (∧,∨, ¬), where an *atom* is a predicate which valuates to True or False given an assignment to the variables. A *literal* is either an atom or its negation. A *theory* gives special meanings, known as *interpretations*, to functions and predicate symbols within the theory. In this paper, we consider only *the theory of Integer Difference Logic (IDL)*, which requires atoms to be of the form *x*_*i*_ *− x*_*j*_ *≤ c*, where *x*_*i*_ and *x*_*j*_ are integer variables, *c* an integer constant, “ − ” the integer subtraction function, and “ ≤ ” the usual binary ordering predicate. A theory solver decides the satisfiability of a conjunction of *literals*. In particular, an IDL theory solver can be implemented as the Bellman-Ford algorithm which runs in polynomial time. Incorporating a SAT solver and a theory solver, an SMT solver takes in a formula and outputs an assignment to the variables if the formula is satisfiable or otherwise reports unsatisfiability.

We observed that SMT is a natural way of encoding the Wheeler graph recognition problem. Firstly, for each node a variable is created representing the ordering of the node. Recalling the constraints, for all pairs of edges (*u, v*) and (*u′, v′*) labeled *a* and *a′* respectively: (i) *a* ≺ *a*′ → *v* < *v*′, and (ii) *a* = *a*′ ∧ *u* < *u*′ → *v* ≤ *v*′. By indexing the known labels lexicographically, we obtained an SMT formula containing constraints (i) and (ii), in which all atoms are of the form *v* ≤ *v*′ satisfying the IDL requirement. Note that the strict inequality *v* < *v*′ can be rewritten using the the non-strict one as *v* − *v*′ ≤ 1. Besides constraints (i) and (ii), we also enforced all-different constraints and range constraints for all nodes, which have the form 1 *≤ v ≤ n*. The node orderings can be obtained from the satisfying assignment by solving the SMT formula iff it is satisfiable. In `Wheelie-SMT`, we used Z3 [16] as the underlying SMT solver.

As the number of constraints and variables are the main factors affecting runtime, simplifying the problem or providing additional information can improve performance. We noticed that the problem can be simplified using the rough order obtained from the *renaming heuristic* described in Section 3.1. By adding the range information to the formula, notice that constraint (i) can be removed from the formula. Moreover, the all-different constraints of nodes from different groups can also be removed. The rationale is that if the graph was not reported non-Wheeler during the *renaming heuristic*, the range constraints provided by the procedure must automatically imply constraint (i). The benefits are twofold: not only the number of constraints is reduced, the search space is also significantly pruned.

### 3.3 WGT’s graph generating algorithms

We implemented five generators in Python scripts to produce tries, reverse deterministic graphs [5], De Bruijn graphs and *complete* and *d-NFA* random Wheeler graphs. The first three generators take either DNA or protein multiple sequence alignments and produce the corresponding graph structures. As for the two random generators, users can produce a Wheeler graph given its *n, e*, and *σ*, and *d-NFA* random generator can further take *d*, which controls the most number of edges coming out from a node with the same label (*d*-NFA) as user input.

Tries and De Bruijn graphs are Wheeler graphs by definition. These generators start by removing gap placeholders from the multiple alignments. The trie generator iterates through the prefixes of each sequence in the multiple alignment, inserts characters into the trie and creates a new node at the end of a path if a prefix cannot be traversed from the source. Edges are labelled according to the label of the parent. The De Bruijn graph generator constructs a distinct *k −* 1-mer dictionary from the sequences. It connects edges between adjacent two nodes and label the edge with the first character in the *k −* 1-mer of the child node. Reverse deterministic graphs are usually invalid Wheeler graphs but might be valid when the graphs are small, and once violations occur, adding more nodes and edges cannot turn them back to Wheeler graphs. The reverse deterministic graph generator iterates through columns of a multiple sequence alignments from right to left. At a column *i*, it creates distinct nodes for the characters found there, connecting them to the current node with the node of the previous ungapped character with the direction pointing to the end of the alignments and the label of the previous ungapped character. This follows the procedure described in the GCSA study [5]. Last, three generators initializes the names of nodes with the breadth first search orders and outputs the constructed graph in DOT format.

#### Algorithm 1

Wheeler graph Recognition Algorithm, `Wheelie`

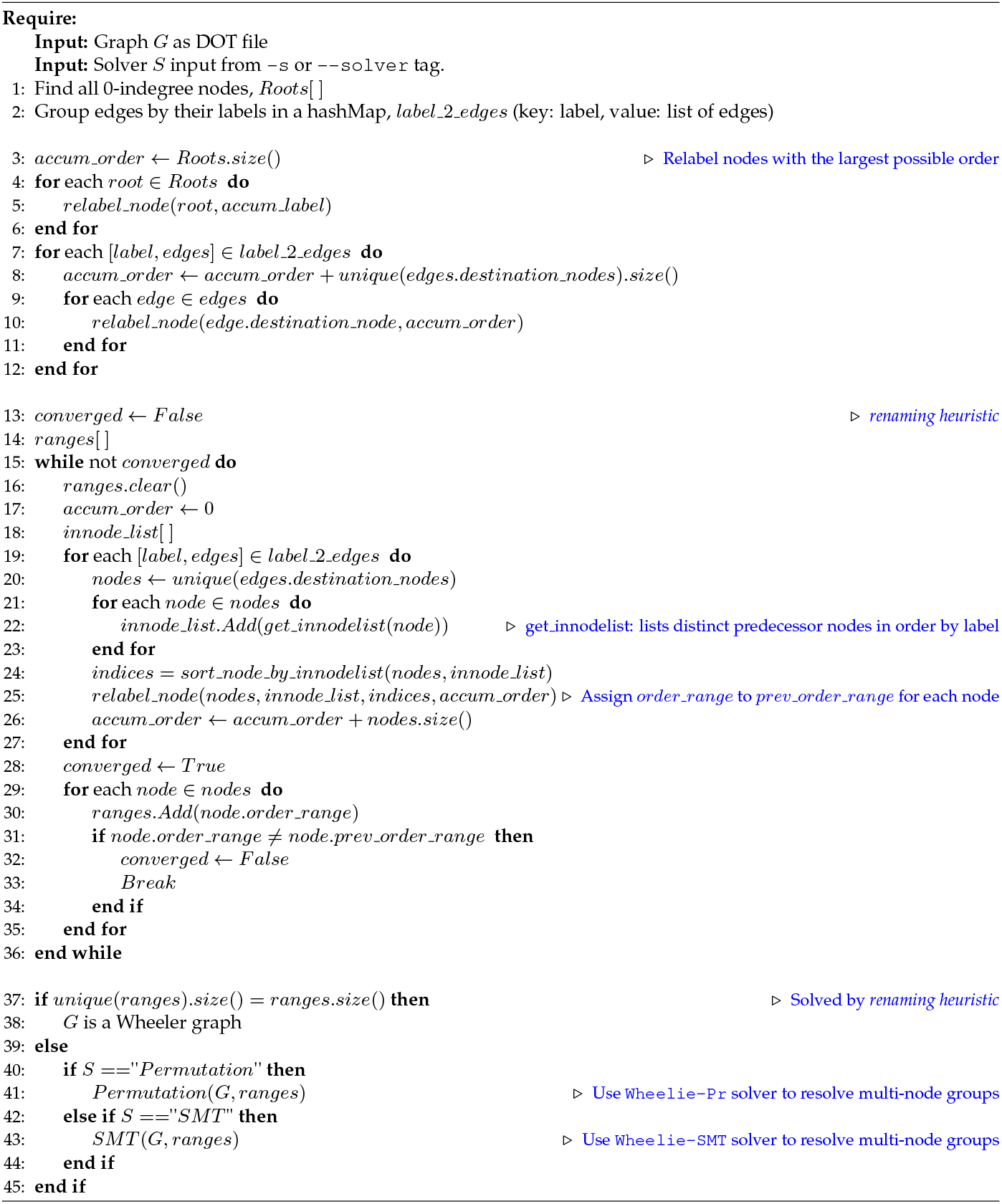

For the two random generators, *complete* Wheeler graph and *d-NFA* Wheeler graph generators, we explain our design concepts in Appendix section2.

## 4 Discussion

We demonstrated that `Wheelie-SMT` is the fastest and most robust algorithm available for the Wheeler Graph recognition problem. We showed this across a variety of graph types, including large graphs (thousands of nodes and edges) and challenging graphs, such as those that are *d*-NFAs with values of *d* up to 8. We also demonstrated WGT’s facilities for visualizing and understanding these graphs.

While current pan-genome representations tend to be based on De Bruijn graphs, which are Wheeler Graphs, other relevant pan-genome graph representations are not necessarily Wheeler. For example, the reverse deterministic automata of the GCSA study [5] are not Wheeler, though they can be made Wheeler through a “path doubling” process. In the future, we expect that WGT and the Wheelie algorithm will be useful for studying alternative pan-genome graph representations that might improve upon De Bruijn graphs in various ways. In Appendix Figure 3, we provide a simple illustration of why another Wheeler Graph (besides a De Bruijn graph) might be better suited as a pan-genome representation, since it (a) uses fewer nodes and edges than a corresponding De Bruijn graph, (b) does not require that we select a particular value for *k* (the *k*-mer length), and (c) avoids collapsing sequences that are distinct with respect to the coordinate system of a given genome.

We noted a relationship between the renaming heuristic proposed here and the “forward algorithm” of Alanko et al [15]. In the future, it will be important to clarify the relationship between these two algorithms, which have similar goals but take different approaches to partitioning and ordering the graph nodes. Appealingly, the forward algorithm has polynomial running time when the input is already a Wheeler graph; we do not have a similar guarantee for the renaming heuristic. However, the forward algorithm is not directly usable for the same purpose as the renaming heuristic, since it is possible for the forward algorithm to collapse a non-Wheeler input in a way that produces a Wheeler output. Another question for future work is whether the renaming heuristic could be combined with the forward algorithm to obtain an algorithm with strong guarantees (like the forward algorithm) but that is directly applicable to the recognition problem (like the renaming heuristic).

When `Wheelie` determines that a graph is is Wheeler Graph, it is able to report a node ordering that can then be used to index the graph. In the future, it will be important to extend `Wheelie` to report other useful information, including when the graph is not a Wheeler Graph. For instance, when `Wheelie` encounters a conflict that proves the graph to be non-Wheeler, `Wheelie` could supply the user with an explanation for why the graph cannot be Wheeler. Such an explanation could also allow `Wheelie` to suggest modifications to the graph that would make it a Wheeler Graph, without changing which strings it encodes. A trivial example would be a node with two incoming edges having two distinct labels. This violates the Wheeler Graph properties, but also suggests a potential solution: the node could be duplicated, with outgoing edges also duplicated. The initial inbound edges could be redrawn to point to the distinct duplicates, possibly restoring the Wheeler properties. A more general approach for understanding Wheeler violations could work by extracting conflicting sets of clauses from the SMT algorithm, and converting them into a human-understandable or other actionable form.

It may also be possible to encode the *renaming heuristic* as a set of clauses in the SMT solver, potentially allowing the entire algorithm to execute within the SMT solver. Finally, as different SMT solvers such as CVC5 [17] or Z3 [16] adopt different heuristics, they could potentially be substituted into WGT, or combined for increased efficiency [18].

## Supporting information

Wheeler_Graph_Toolkit_appendix.pdf

## 5 Acknowledgments

We thank Markus J. Sommer for proposing the name `Wheelie`. We thank Nicola Prezza, Nicola Cotumaccio, Travis Gagie and Christina Boucher for helpful comments.

## 7 Authors’ contributions

KC, PC and BL designed the method. KC and PC wrote the software and performed the experiments. KC, PC, SAS and BL wrote the manuscript. All authors read and approved the final manuscript.

