## Supplementary material for "WGT: Tools and algorithms for recognizing, visualizing and generating Wheeler graphs": Wheeler_Graph_Toolkit_appendix.pdf

April 12, 2023

### 1 `Wheelie`'s permutation based approach

While the G & T algorithm explores an exponential-sized space of possible array assignments, `Wheelie` explores a factorial-sized space of node permutations. This may or may not lead to a larger search space for `Wheelie`, depending on the graph's properties. To be specific, the G & T's algorithm may have to consider all $2^{2(e+n)+e\log(\sigma)}$ assignments for $I, O$ and $L$. Our approach might need to consider $n!$ node permutations in the worst case. We sought a rough comparison between the approaches in light of the fact that G & T's space depends not only on $n$ but also on $e$ and $\sigma$. Below Derivation 1, we fixed $n$ for both, defining a new variable $C$ as $e(2+\log\sigma)$. We then found some values for $C$ that equalize the algorithms' search space size under various values for of $n$s (Table 1). For instance, when $n = 100$, $C$ can be at most 324 in order for G & T's algorithm has an equal or smaller search space than `Wheelie-Pr`, which is a strict threshold, and furthermore, this comparison is done with `Wheelie-Pr` skipping the *renaming heuristic*, which in reality makes `Wheelie-Pr` superiorly faster (Results2.4).

To gain a further advantage over the G & T algorithm, `Wheelie` further strives to prune the search space, using a renaming heuristic, an SMT solver, or both, as detailed in Methods3.

$$2^{2(n+e)+e\times\log\sigma} = n! \quad (1)$$

$$2^{2n+e\times(2+\log\sigma)} = n! \quad (2)$$

$$\text{Let } C := e(2+\log\sigma)$$

$$2n + C = \log_2 n! \quad (3)$$

$$C = \sum_{x=1}^{n} \log_2 x - 2n \quad (4)$$

Derivation 1: The relationship between $C$ and $n$

| n | $C$ threshold |
|---|---|
| 10 | 1.79 |
| 20 | 21.08 |
| 30 | 47.71 |
| 40 | 79.16 |
| 60 | 152.13 |
| 80 | 234.83 |
| 100 | 324.76 |
| 150 | 572.86 |
| 200 | 845.38 |

**Table 1:** Threshold values of $C$ as a function of $n$ such that values greater than the threshold cause the permutation-based approach to have a smaller search space compared to the G & T approach.

### 2 Random generators

We implemented two random generators, a *complete* Wheeler graph generator and a *d-NFA* Wheeler graph generator. We first fix the ordering of nodes and then try to select edges such that both user-specified constraints and Wheeler graph properties are satisfied. Let $N_i$ be the nodes with incoming edges labeled $i$ and $E_i$ be the edges labeled $i$ where $i = 1, 2, ..., \sigma$, and also let $n_i = |N_i|$ and $e_i = |E_i|$. In both generated graphs, we assume that $n_i \approx \frac{n-r}{\sigma}$ and $e_i \approx \frac{e}{\sigma}$ where $r$ is the number of nodes without incoming edges.

We say a Wheeler graph $G$ is *complete* if no more edges can be added to $G$ while maintaining the Wheeler graph properties.

**Property 1.** *Given number of nodes $n$ and number of labels $\sigma$, the number of edges of a Wheeler graph is upper bounded by $e_{max}$ where*

$$e_{max} = n \times \sigma + n - \sigma - r = (n-1)(\sigma+1) - r + 1 \tag{5}$$

*Proof.* Consider the bipartite representation of a Wheeler graph $G$ with number of nodes $n$ and number of labels $\sigma$. Note that

$$\sum_{i=1}^{\sigma} n_i = n - r. \tag{6}$$

Observe that for each label $i$, the number of edges that is labeled $i$ is at most $n + n_i - 1$. Taking the sum of edges of each label and applying Equation (6), we have

$$e_{max} = \sum_{i=1}^{\sigma}(n + n_i - 1) = n \times \sigma - \sigma + \sum_{i=1}^{\sigma} n_i = n \times \sigma - \sigma + n - r. \tag{7}$$

□

One way of generating complete Wheeler graphs is to have all nodes connect to the first node of $N_i$ and the last node additionally connect to the rest of the nodes in $N_i$ for each label $i$ (the last node has $n_i$ outgoing edges in total). By randomly selecting $n_i - 1$ nodes from $N$ and connecting the selected nodes to consecutive nodes in $N_i$, a new complete Wheeler graph can be generated by appropriately shifting the destination node of each edge such that the Wheeler graph property is maintained. Figure 3 shows an example of a complete Wheeler graph with $(n, e, \sigma, r) = (7, 18, 2, 1)$. With a complete Wheeler graph of $n$ nodes and $\sigma$ labels, we are able to generate random Wheeler graphs with $e < e_{max}$ edges by sampling $e$ distinct edges from the complete Wheeler graph.

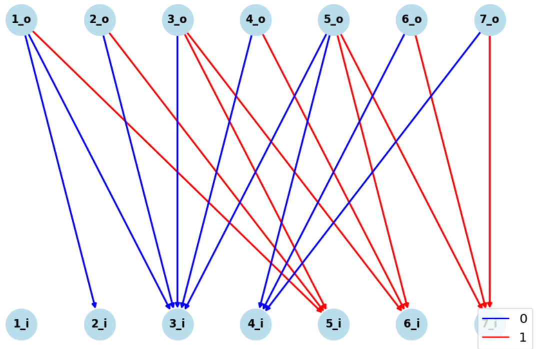


**Figure 1:** A complete Wheeler graph with $(n, e, \sigma, r) = (7, 18, 2, 1)$. In this example we have $n_0 = n_1 = 3$. The selected nodes for label $0$ is node 1 and 5 and for label $1$ node 3 and 5.

For generation of d-NFA Wheeler graphs, let $x_k$ be the number of nodes with $k$ outgoing edges of the same label $i$. Thus, given $e$ and $\sigma$ we have

$$\sum_{k=1}^{d} k \cdot x_k = e_i \tag{8}$$

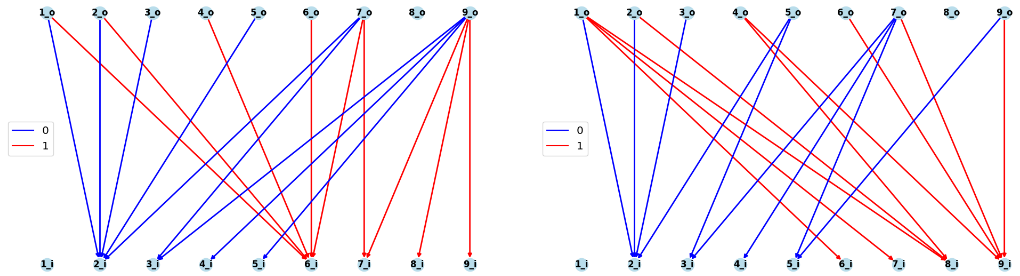


**Figure 2:** A 3-NFA Wheeler graph with $(n, e, \sigma, r) = (9, 18, 2, 1)$. On the left shows the original Wheeler graph and on the right shows the Wheeler graph after swapping nodes and corresponding edges. Our d-NFA generator directly generates Wheeler graphs with swapped nodes and edges.

Also the number of nodes $n_i$ with incoming edges labeled $i$ must be greater than the minimum number of nodes $n_{min}$ needed to accommodate $e_i$ edges. Thus, we have

$$n_{min} = 1 + \sum_{k=2}^{d} (k-1) \cdot x_k \leq n_i \tag{9}$$

Note that any solution $x_k$ that satisfies Equation (8) and (9) for all edge label $i$ represents a set of valid Wheeler graphs. To see this first notice that given $x_k$, we can always order the nodes such that $x_1$ nodes with one outgoing edge are placed at the front, followed by $x_2$ nodes with two outgoing edges, followed by $x_3$ nodes with three outgoing edges and so on. By construction, this gives a valid d-NFA Wheeler graph. Moreover, by swapping the nodes and reconnecting the corresponding edges accordingly, different d-NFA Wheeler graphs can be obtained. An example is shown in Figure 2.

To obtain a concrete instance, we find a valid solution for $x_k$ and then determine the node ordering. In our case, we set all $x_k$ to be the same, and if not possible assign the residual to $x_1$ to satisfy Equation (8). We believe that this reflects the hardness of different benchmarks with different d's. For node ordering, all the nodes are shuffled and edges are distributed such that each node in $N_i$ has at least one incoming edge while maintaining the Wheeler properties.

#### 3 The benefit of recognition Wheeler graphs from reverse deterministic graphs

Figure 3 shows an example of creating a De Bruijn graph and a reverse-deterministic graph for a multiple sequence alignment.

The workflow on the left-hand side shows the process of creating a k=4 De Bruijn graph. In the first step, gaps in the multiple sequence alignment are removed, and sequences are chopped into 3-mers (step 2). In step 3, nodes with the same 3-mers are merged, and the first character of each k-mer node is labelled on the edge that goes into it (step 4). In step 5, we do breadth-first search to relabel k-mer nodes. Finally, a De Bruijn graph with 18 nodes and 21 edges is created.

The process of creating a reverse deterministic graph is illustrated in the workflow on the right-hand side. In Step 1, we add ′ at the beginning and ′$ at the end of sequences. Step 2 involves merging nodes in each column that share the same sequences. These merged nodes are labeled with the node where the edge goes into, as shown in Step 3. Next, we relabel the nodes using breadth-first search in Step 4. If the current graph is not a Wheeler graph, as checked in Step 5, we unzip the nodes into bubbles. For instance, in this example, node 9 can be unzipped into node 9 and node 9′, and node 6 can be unzipped into node 6 and node 6′, resulting in a Wheeler graph with 14 nodes and 16 edges. Finally, we can run our Wheeler Graph recognizer to index both the De Bruijn Graph and the reverse deterministic graph in step 6.

There are two main benefits of modifying a reverse deterministic graph into a Wheeler graph over simply indexing a genome into a De Bruijn graph. The first reason is that in some cases, it is possible to create a smaller graph. Figure 3 is an example. The final Wheeler Graph created from the reverse-deterministic graph, which has 14 nodes and 16 edges, is more compact compared to the k=4 De Bruijn graph, which has 18 nodes and 21 edges. The second reason is that the reverse deterministic graph maintains the original coordinates after collapsing states from a multiple sequence alignment. As shown in Step 3 in Figure 3, nodes labelled with the same k-mers are merged into the same nodes. There are some k-mer nodes far away in different positions in the genome that are merged, which messes up the coordinates. It makes the downstream pattern-matching process harder.

Based on the two reasons above, we can see the advantages of indexing a graph into a Wheeler graph. Its potential application is to fix reverse deterministic graphs, which are not always indexable, into Wheeler graphs, and it can be applied to build the pangenome.

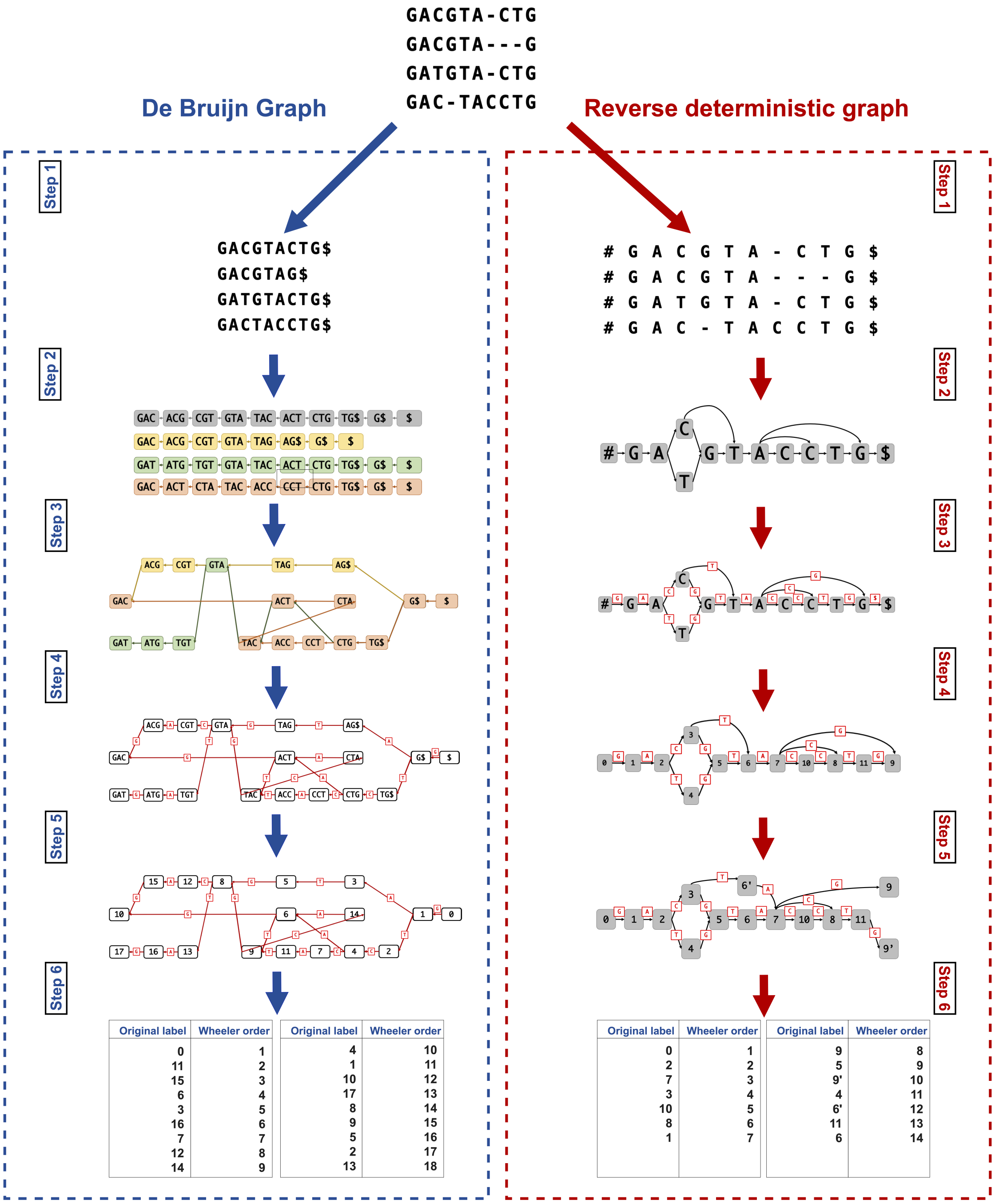


| Original label | Wheeler order | Original label | Wheeler order |
|---|---|---|---|
| 0 | 1 | 4 | 10 |
| 11 | 2 | 1 | 11 |
| 15 | 3 | 10 | 12 |
| 6 | 4 | 17 | 13 |
| 3 | 5 | 8 | 14 |
| 16 | 6 | 9 | 15 |
| 7 | 7 | 5 | 16 |
| 12 | 8 | 2 | 17 |
| 14 | 9 | 13 | 18 |

| Original label | Wheeler order | Original label | Wheeler order |
|---|---|---|---|
| 0 | 1 | 9 | 8 |
| 2 | 2 | 5 | 9 |
| 7 | 3 | 9' | 10 |
| 3 | 4 | 4 | 11 |
| 10 | 5 | 6' | 12 |
| 8 | 6 | 11 | 13 |
| 1 | 7 | 6 | 14 |

**Figure 3:** The example workflow of creating a De Bruijn graph and a reverse-deterministic graph.
